# Evaluating Cost-Effective Methods for Rapid and Repeatable National Scale Detection and Mapping of Invasive Species Spread

**DOI:** 10.1101/353722

**Authors:** Ruth A. Kost, Ryan K. Brook

## Abstract

Invasive species can spread rapidly at local and national scales, creating significant environmental and economic impacts. A central problem in mitigation efforts is identifying methods that can rapidly detect invasive species in a cost-effective and repeatable manner. This challenge is particularly acute for species that can spread over large areas (>1 million km^2^). Wild pigs (*Sus scrofa*) are one of the most prolific invasive mammals on Earth and cause extensive damage to agricultural crops, native ecosystems, and livestock, and are reservoirs of disease. They have spread from their native range in Eurasia and North Africa into large areas of Australia, Africa, South America, and North America. We show that the range of invasive wild pigs has increased exponentially in Canada over the last 30 years following initial and ongoing releases and escapes from domestic wild boar farms. We evaluate eight different methods for mapping invasive species over large areas and assess their benefits and limitations. Our findings effectively map the spread of a highly invasive large mammal and demonstrate that management efforts should ideally rely on a set of complementary independent monitoring methods. Mapping and evaluating resulting species occurrences provide baseline maps against which future changes can be rapidly evaluated.

## Introduction

The Anthropocene is the current geological age on earth, characterized by the dominant influence of humans on the environment, including impacts on native ecosystems and spread of invasive species globally^1^. Indeed, invasive species have been identified as one of the greatest threats to global biodiversity^2^. Human activities can facilitate the spread of invasive species, directly through intentional and unintentional movements of animals, and indirectly through habitat fragmentation and change associated with agriculture and urban expansion, climate change, and over harvest of native species^3,4,5^. Efforts to mitigate the spread of invasive species have been limited by the lack of timely and accurate maps of occurrences and spatial expansion, especially over very large areas (>1 million km^2^). Requirements for national scale mapping are limited by project budgets and new approaches are needed that are cost-effective and repeatable, especially for species expected to expand rapidly over large areas and that are associated with high potential for harm.

Wild pigs (*Sus scrofa*), also referred to as feral swine, wild hogs, or feral hogs^6^, currently have the largest global range of any non-domesticated terrestrial mammal on earth^7^. Native to Eurasia and part of North Africa, wild pigs have expanded their range, primarily through human introductions combined with natural dispersal, across all continents except Antarctica^7,8^. The broad geographic extent of their native range, coupled with the generalist nature of the species has allowed wild pigs to easily adapt and survive in new environments that span a broad range of climate, habitat, and resources^9,10^. The widespread success of wild pigs is explained by their extremely high fecundity^11^, early sexual maturity^12^, plastic diet^7^, long lifespans^13^, and highly adaptive nature^9^.

Wild pigs are an invasive species in North America and are descendants of Eurasian wild boar (*S. scrofa scrofa*), domestic pigs (*S. scrofa domesticus*), and hybrids of the two^14,15^. Long-established populations have existed in southern parts of the U.S. for hundreds of years, with areas of high population densities in Texas, Florida, and California. In the continental United States, there has been a well-documented expansion in the distribution and abundance of wild pigs in recent decades, from 17 to 38 states during the last 30 years^16^. Wild pigs were first introduced to Canada during a federal and provincial agriculture diversification initiative in the 1980’s and 1990’s to diversify livestock species and supplement producer incomes^17,18^. Escapes and intentional releases from domestic wild boar farms have led to the feral populations that are established on the Canadian Prairies^18,19^. Brook and van Beest^18^ provided a coarse-scale distribution of wild pigs in Saskatchewan at the Rural Municipality level. However, prior to this current study there has not been a comprehensive national scale map of the species range in Canada.

Within their introduced range, wild pigs pose many environmental and socio-economic threats including, but not limited to; environmental and water quality degradation^20,7^, predation of native wildlife^16^, disease hosts and vectors^21^, damage to agricultural crops and livestock ^22,23^, and threats to human safety^24^. Of particular concern is the threat wild pigs pose to Canada’s ecosystems and the native wildlife and species at risk that inhabit them, as well as the significant economic losses that could be incurred to the livestock industry and international trade if reportable diseases were identified^21,25^.

Wild pigs are known hosts and reservoirs of numerous viral and bacterial diseases, as well as parasites, which are infectious to wildlife, livestock, and humans^21,16,26^. Disease threats are an increasing concern to livestock producers and the pork and beef industries, as disease outbreaks are associated with high economic losses^26,9^. Common diseases of concern transmitted from wild pigs to livestock are swine brucellosis, bovine tuberculosis, pseudorabies, and classical swine fever, while zoonotic diseases include brucellosis, *Escherichia coli*, salmonellosis, and leptospirosis ^16,21,27,28^. Negative impacts associated with wild pigs have been well-documented across Europe, Australia, and the United States; however, have not been documented in Canada. Ecological damage to native vegetation, riparian areas, water quality, wildlife habitat, and protected or endangered species is a large concern associated with the presence of wild pigs^8^. Agriculture losses in the United States from wild pig damage have yet to be fully quantified, but an estimate of $1.5 billion USD per year was presented by Pimental^29^ based on extrapolation of localized data from some areas. Wild pig-vehicle collision cost $36 million USD per year^30^. There has been no research to characterize economic, social, or environmental impacts in Canada.

A key challenge in managing rapidly expanding invasive species such as wild pigs at national and continental scales is having up-to-date information on their spatial distribution. Mapping the locations of invasive species is central to guiding effective management and is essential to determine if control efforts are effective at controlling and limiting, or even reducing, their spatial expansion^31^. However, identifying cost-effective methods to accurately and repeatedly map a species at a national scale represents a significant time and financial commitment. Conventional ecological monitoring used for large mammals such as aerial surveys, trail cameras, and mark-recapture can be effective at relatively small scales (<100,000km^2^), but become time and cost prohibitive at much larger scales. The use of local and traditional knowledge accumulated by people living and working on the land through personal observations and shared knowledge has been used for documenting species occurrences^32,33^. Such data can be collected using personal interviews, mail surveys, internet surveys, telephone surveys, and open source mapping^34,35^. Similarly, citizen science engages large numbers of lay people in collecting species occurrence data, such as the annual Breeding Bird Count and the Christmas Bird Count across the United States and Canada^36^, though this also requires considerable logistics coordination and has rarely been used for large mammals. Efforts to incorporate local knowledge in data collection using rigorous social science methods have rarely been applied over large areas (>1 million km^2^). Much work remains to evaluate the efficacy of these methods and determine the benefits and limitations of the different approaches in the face of immediate needs for detailed information on invasive species in general and wild pigs in Canada specifically. As such, the objectives of this study were to: (1) identify the current spatial distribution of wild pigs across Canada, and (2) evaluate the benefits and limitations of eight different independent approaches for national-scale, repeatable mapping of wild pig spatial distribution.

## RESULTS

Data were collected from eight independent data methods and five of these (stakeholder snowball sampling, expert interviews, bounty, telephone survey, and GPS collars) were used directly to develop maps of wild pig distribution across Canada using all datasets combined for three time periods, 1990-2000, 2001-2010, and 2011-2017 (Fig.1). A total of 625 occupied watersheds were identified, out of 37,578 watersheds in the study area (1.6% occupied by wild pigs), based on pooled results of all methods across all years. The area of watersheds occupied in Canada has increased exponentially from 1995 to 2017 (Fig. 2). The large majority of the spatial expansion (92%) occurred in the three prairie provinces of Alberta, Saskatchewan, and Manitoba (Fig. 3). Indeed 58% of the national spread of wild pigs occurred just in Saskatchewan (Fig. 3). Wild pigs are also established in localized populations in British Columbia, Ontario, and Quebec. Of the ten provinces in Canada, only the four eastern provinces in Atlantic Canada (Newfoundland and Labrador, New Brunswick, Prince Edward Island, and Nova Scotia; 5% of Canada combined) have no confirmed sightings of wild pigs.

**Figure 1.**
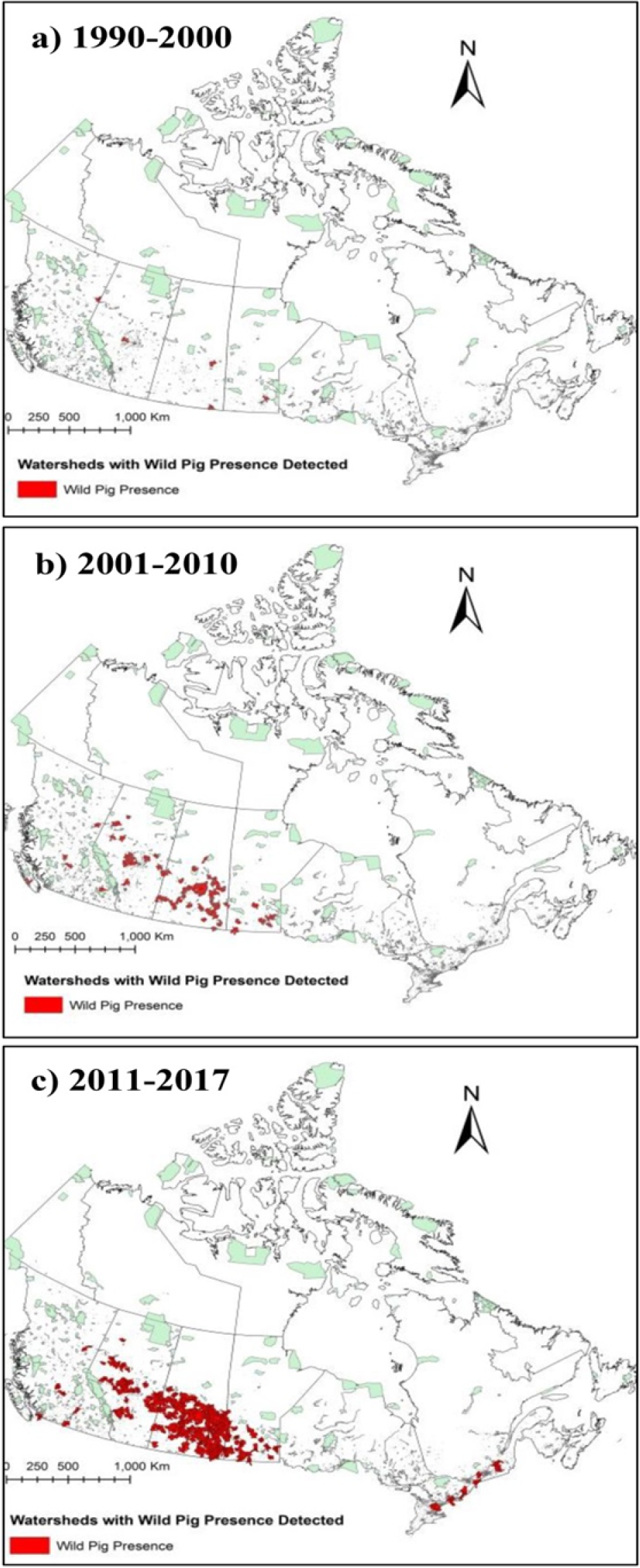
Spatial expansion of wild pigs in Canada from the period of initial escapes and releases (1990) to present, based on combined results from point occurrences obtained from national scale stakeholder interviews, expert interviews, a telephone survey, and locations from a bounty conducted in Alberta mapped. Watersheds where wild pigs were detected are mapped for three time periods, a) 1990-2000, b) 2001-2010, and c) 2011-2017.

**Figure 2.**
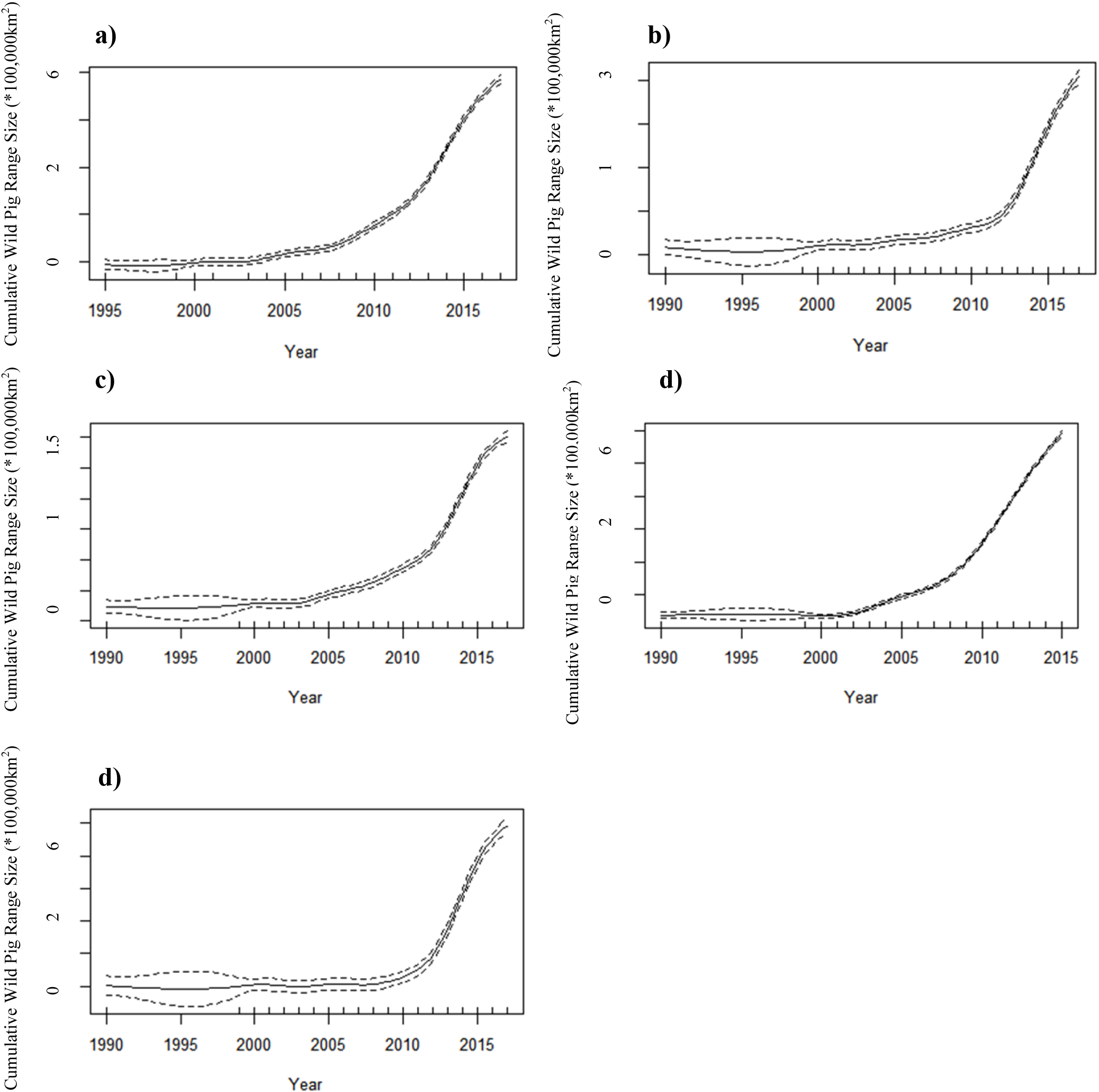
a) Cumulative range size of wild pigs in Canada mapped by occupied watersheds (1990-2017) using a) combined results from point occurrences obtained from all methods; b) stakeholder interviews; c) national scale expert interviews; d) Alberta bounty; and e) national scale rural telephone survey. Results were modelled using a generalized additive model in R (n =19, R^2^= 0.998, p < 0.001) along with a 95% confidence interval.

**Figure 3.**
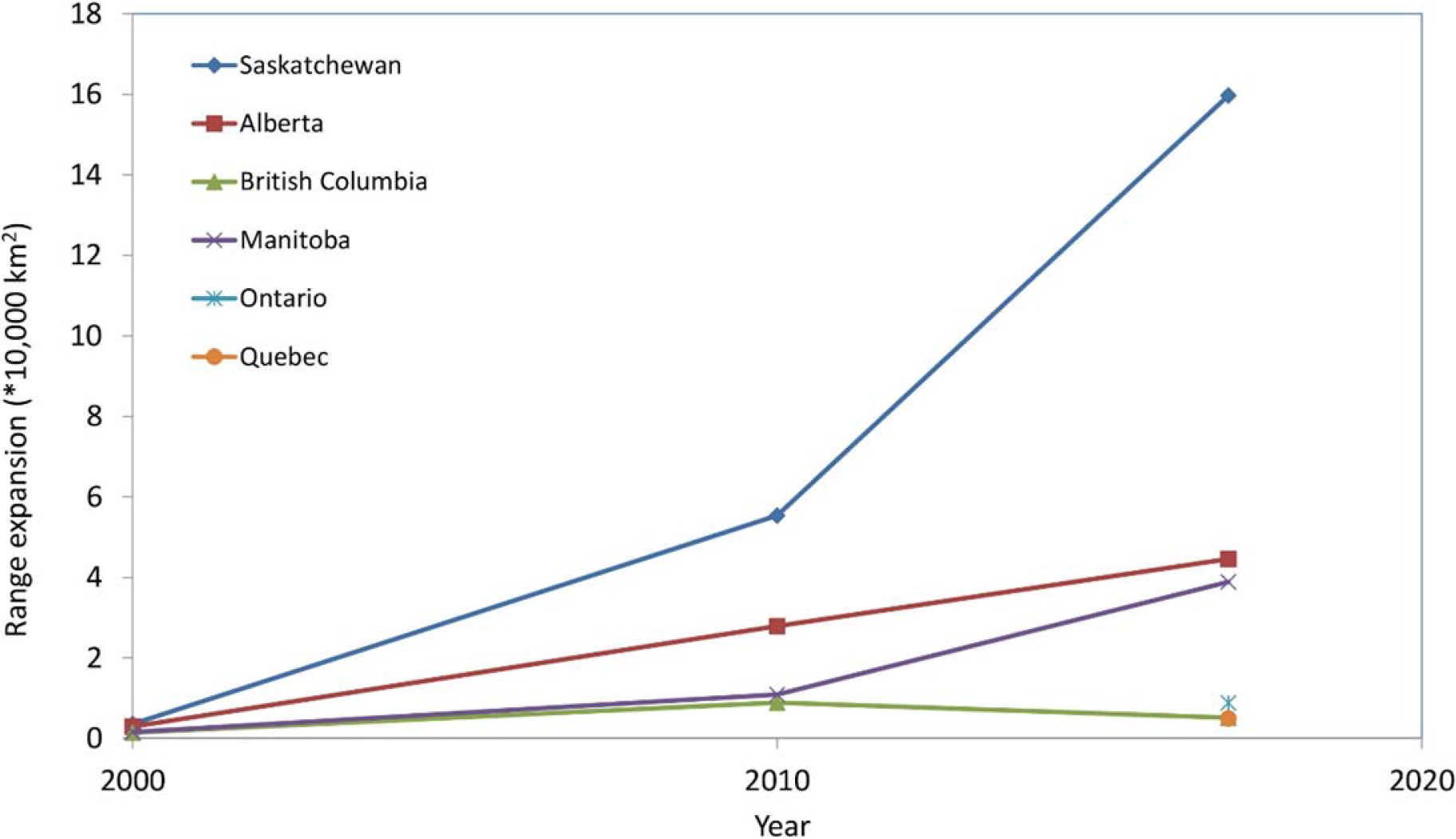
Increase per decade in range size of wild pigs in each of the seven Canadian provinces confirmed to have wild pigs mapped by occupied watersheds (1990-2017) using combined results from point occurrences obtained from national scale stakeholder interviews, expert interviews, a telephone survey, and locations from a bounty conducted in Alberta.

Based on analysis at the level 9 watershed, wild pigs are present in a total area of 262,043 km^2^. The average annual cumulative increase in wild pig range was 5,504 km^2^ from 1990-2017. Over the period 2000-2017 the average provincial increase in wild pig range was 24,941 km^2^.

As expected, the number of watersheds in Canada occupied by wild pigs was different among sampling methods (Table 1). There was an overall high consistency in the spatial distribution of occupied watersheds for each method. However, specific correspondence of individual occupied watersheds between data collection methods was low overall (<50%) for all methods (Table 2). Correspondence was greatest between the occupied watersheds determined using expert interviews and stakeholder snowball interview methods (14% correspondence). The number of occupied watersheds was significantly and positively associated with the level of correspondence among all possible pairs of data collection methods (R^2^ = 0.90, df = 5, p < 0.001).

**Table 1.**
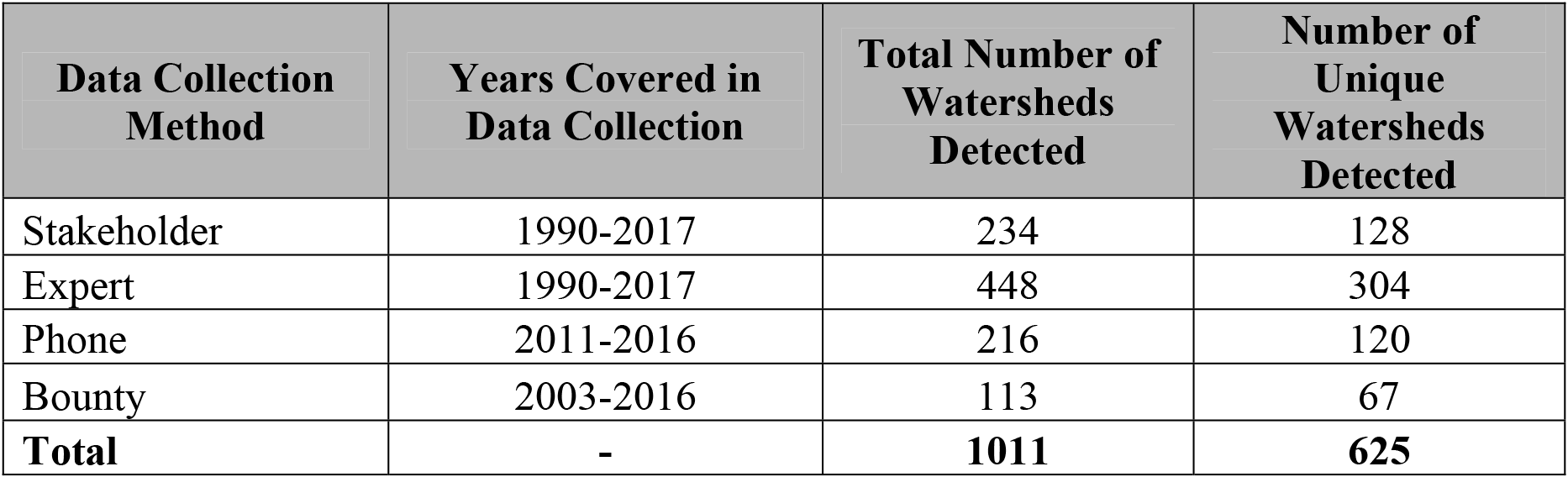
Comparison of the number and distribution of watersheds in Canada with wild pig occurrences, based on four unique monitoring strategies.

**Table 2.** Correspondence of wild pig detection by watershed among different data collection methods to identify wild pig occurrences by watershed in Canada (1990-2017).

|  | Number of Watersheds With Wild Pigs Detected | % Correspondence |
| --- | --- | --- |
| Stakeholder-Expert | 96 | 14 |
| Stakeholder-Phone | 31 | 7 |
| Stakeholder-Bounty* | 12 | 3 |
| Expert-Phone | 62 | 9 |
| Expert-Bounty* | 10 | 2 |
| Phone-Bounty* | 32 | 10 |
| Stakeholder-ALL | 234 | 23 |
| Expert-ALL | 448 | 44 |
| Phone-ALL | 216 | 21 |
| Bounty-ALL | 113 | 11 |

\*Only Alberta locations were used for correspondence with bounty as this is the only province that implemented a bounty during this study.

Overall mean response rate for the expert interviews was 51% (S.E. = 5.9), with a standard error of 5.89. The mean number of completed districts within provinces was 55% (S.E. = 7.7). Total number of individual participants in the stakeholder snowball sampling study was 275.

Each of the eight research methods evaluated to determine wild pig distribution had important benefits and limitations and were viewed by stakeholders and other participants with varying degrees of creditability and spatial accuracy (Table 3). Of the social science methods used, snowball sampling produced the most wild pig occurrences. GPS collared pigs produced an extremely large number of accurate locations from a small number of animals over 4-25 months. The three social methods (expert interviews, stakeholder snowball sampling, and telephone survey) resulted in the majority (86%) of all watersheds detected with wild pigs in this study. Total cost of data collection for all eight methods used was $918,416 over the seven years of this study. Average cost per wild pig occurrence by data collection method was $336 (range $4 -$1,291). Average cost per watershed detected with wild pigs present was $6,540 (range $93 -$45,643).

**Table 3.**
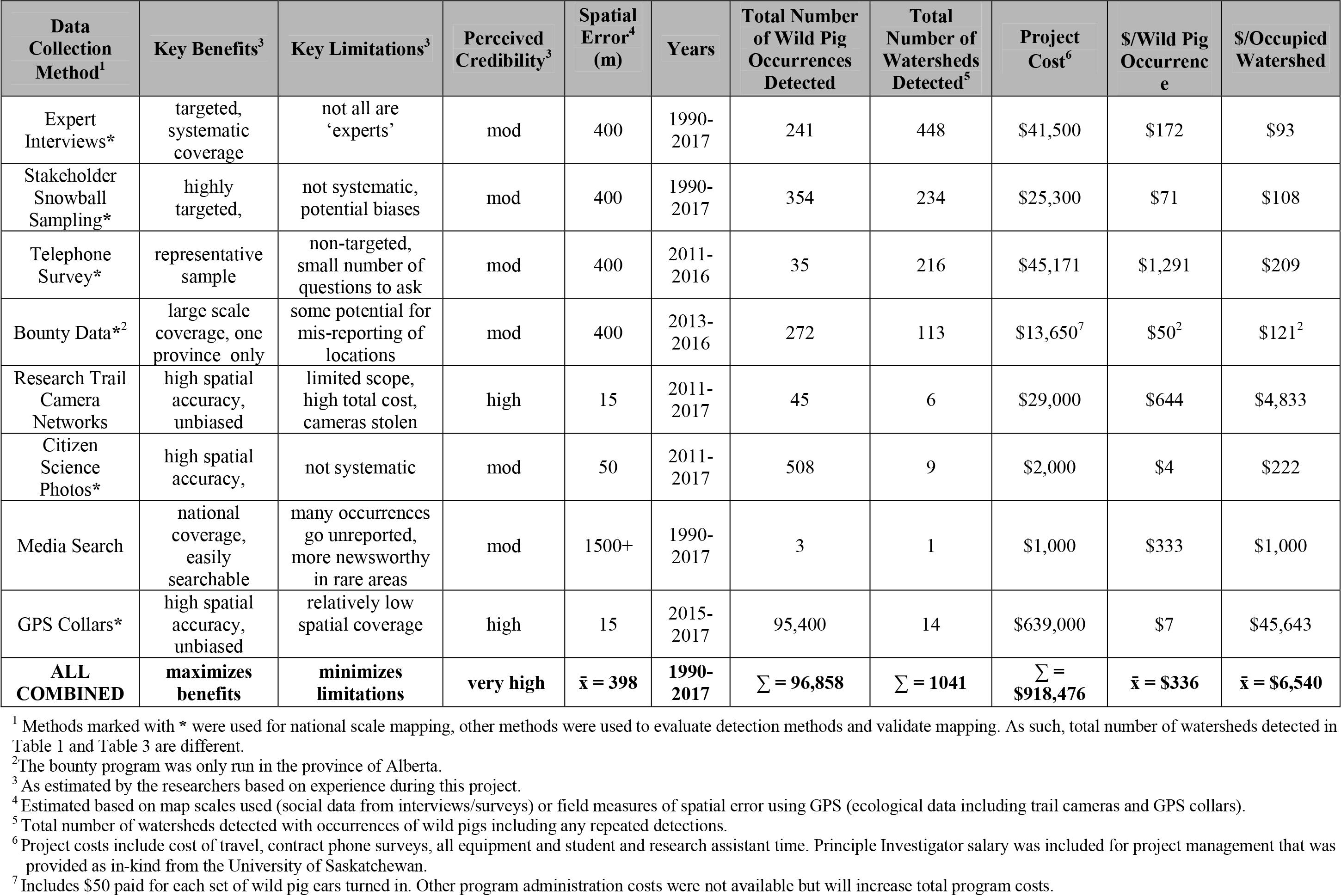
Summary of financial costs, benefits, and limitations of eight different methods for detecting invasive wild pig occurrences and occupied watersheds across Canada collected during 2011-2017 that cover the time period 1990-2017.

| Data Collection Method<sup>1</sup> | Key Benefits<sup>3</sup> | Key Limitations<sup>3</sup> | Perceived Credibility<sup>3</sup> | Spatial Error<sup>4</sup> (m) | Years | Total Number of Wild Pig Occurrences Detected | Total Number of Watersheds Detected<sup>5</sup> | Project Cost<sup>6</sup> | \$/Wild Pig Occurrence | \$/Occupied Watershed |
| --- | --- | --- | --- | --- | --- | --- | --- | --- | --- | --- |
| Expert Interviews* | targeted, systematic coverage | not all are 'experts' | mod | 400 | 1990-2017 | 241 | 448 | \$41,500 | \$172 | \$93 |
| Stakeholder Snowball Sampling* | highly targeted, | not systematic, potential biases | mod | 400 | 1990-2017 | 354 | 234 | \$25,300 | \$71 | \$108 |
| Telephone Survey* | representative sample | non-targeted, small number of questions to ask | mod | 400 | 2011-2016 | 35 | 216 | \$45,171 | \$1,291 | \$209 |
| Bounty Data*<sup>2</sup> | large scale coverage, one province only | some potential for mis-reporting of locations | mod | 400 | 2013-2016 | 272 | 113 | \$13,650<sup>7</sup> | \$50<sup>2</sup> | \$121<sup>2</sup> |
| Research Trail Camera Networks | high spatial accuracy, unbiased | limited scope, high total cost, cameras stolen | high | 15 | 2011-2017 | 45 | 6 | \$29,000 | \$644 | \$4,833 |
| Citizen Science Photos* | high spatial accuracy, | not systematic | mod | 50 | 2011-2017 | 508 | 9 | \$2,000 | \$4 | \$222 |
| Media Search | national coverage, easily searchable | many occurrences go unreported, more newsworthy in rare areas | mod | 1500+ | 1990-2017 | 3 | 1 | \$1,000 | \$333 | \$1,000 |
| GPS Collars* | high spatial accuracy, unbiased | relatively low spatial coverage | high | 15 | 2015-2017 | 95,400 | 14 | \$639,000 | \$7 | \$45,643 |
| **ALL COMBINED** | **maximizes benefits** | **minimizes limitations** | **very high** | $\bar{x} = 398$ | **1990-2017** | $\Sigma = 96,858$ | $\Sigma = 1041$ | $\Sigma = \$918,476$ | $\bar{x} = \$336$ | $\bar{x} = \$6,540$ |

<sup>1</sup> Methods marked with \* were used for national scale mapping, other methods were used to evaluate detection methods and validate mapping. As such, total number of watersheds detected in Table 1 and Table 3 are different.
<sup>2</sup> The bounty program was only run in the province of Alberta.
<sup>3</sup> As estimated by the researchers based on experience during this project.
<sup>4</sup> Estimated based on map scales used (social data from interviews/surveys) or field measures of spatial error using GPS (ecological data including trail cameras and GPS collars).
<sup>5</sup> Total number of watersheds detected with occurrences of wild pigs including any repeated detections.
<sup>6</sup> Project costs include cost of travel, contract phone surveys, all equipment and student and research assistant time. Principle Investigator salary was included for project management that was provided as in-kind from the University of Saskatchewan.
<sup>7</sup> Includes \$50 paid for each set of wild pig ears turned in. Other program administration costs were not available but will increase total program costs.

## DISCUSSION

Invasive wild pigs are widespread and rapidly expanding their range in Canada, creating important risks to sustainability of native ecosystems and agricultural production following their escapes and purposeful releases from multiple sources at meat and penned shoot farms, starting in 1990. This study has produced the first published maps that characterize the spread of wild pigs across Canada. We show that the majority of the expansion has occurred in the three prairie provinces of Alberta, Manitoba, and especially Saskatchewan. We expect that given the range of available habitats^18^, high reproduction rates with an average of 6 young per female and apparently continuous breeding^37^, and the overall absence of national and provincial management plans and control efforts wild pigs are expected to continue to expand exponentially over the next decade at least. This is consistent with the rapid expansion of wild pig populations in the United States^38,16^ and most other areas of their native and introduced range^39,40,41^. Many areas in Canada are susceptible to wild pig expansion, especially those that are comprised of ample, energy-rich food resources from agriculture crops^10^, forest cover^42^, relatively low predator densities^42^, and repeated introduction/re-introduction events^19^.

Throughout their introduced range, wild pigs pose numerous environmental and socio-economic threats. As an invasive species within Canada wild pigs present a significant threat to Canada’s natural ecosystems and native species. Referred to as ecological train wrecks, wild pigs alter ecosystem processes, vegetation successional stages, nutrient cycles, and cause erosion, sedimentation, and eutrophication to riparian areas and water bodies^20,16,7,43,44,45^. The generalist nature and plastic diet of wild pigs allows them to utilize and compete for a wide variety of resources, as well as predate small mammals, amphibians, invertebrates, and ground nests^16^. The significant disturbance of habitat, resources, and ecosystem processes has direct and indirect effects on native wildlife and has the ability to decrease biodiversity and cause extirpations and extinctions^20,46,47^. Species extirpations and population declines as a result of wild pig presence have been documented in the United States, Galapagos Islands, and Australia^16,48,49,50^. In some areas within wild pig natural and introduced range, populations have expanded to the point where eradication is no longer feasible^51^.

Socio-economic concerns associated with wild pigs are incurred from economic loss from crop damage^52^, predation of small and young livestock^9,44^, vehicle collisions^30^, and structural damage^48^, as well as health and safety concerns to humans, livestock, and wildlife. In many areas of their introduced and native range wild pigs are considered disease reservoirs and maintenance hosts due to their high densities, complex social behaviours, and ability to maintain the disease without a continued source of infection^53,54,55^.

Diseases at the wildlife-livestock interface are multi-faceted and can be transmitted as spill-over or spill-back^56,57^. Diseases at the livestock-wildlife interface pose challenges to wildlife managers and livestock producers, as well as towards quantitative risk analyses as the disease status of wildlife populations and routes of transmission are often poorly understood^21,26^. Disease threats are an increasing concern to livestock producers and the pork and beef industries, as disease outbreaks are associated with high economic losses^9,7,26,56^. The challenges associated with eliminating disease in wildlife populations pose a continued threat of disease introduction to livestock, act as an impediment to disease elimination in livestock populations, and has the potential for spill-over back into wildlife populations ^54,58,59^.

Our finding of wild pigs mainly concentrated in the Prairie Provinces with some of the coldest winters of all ten provinces and that the species is rare or absent in the warm coastal areas is inconsistent with studies that have found the opposite and have found that wild pig distribution is positively associated with warmer climates and have suggested cold winter temperatures to be highly limiting^10,21^. This likely reflects, in part, that domestic wild boar farms that are sources of free-ranging wild pigs were, and are, more concentrated on the Canadian Prairies^19^. The success of wild pigs in western Canada does highlight the capacity for wild pigs to thrive and expand in areas with long and extremely cold winters including some occurrences north of 55° north latitude.

While our mapping efforts provide national scale coverage for an area of 5.5 million km^2^ at a relatively low cost and the benefits of this approach are evident, there are also limitations that should be considered. Importantly, none of the eight methods that were evaluated came close to detecting all of the occupied watersheds by all methods combined. This lack of saturation demonstrates the value of an approach based on multiple methods but also highlights that while expert interviews detected the most wild pig observations within watersheds, they only identify less than half of all watersheds detected in this study. In presence-absence survey methods nondetection errors and false absences are a common sampling problem^60,61^. Due to the elusive and often nocturnal behaviour of wild pigs, their preferred habitat comprised of high levels of hiding cover in wetlands and forests, and relatively low density across most of the study area, some false negatives are expected.

Using interviews with large numbers of study participants and multiple independent methods helped address challenges in detecting wild pigs and balancing the benefits and limitations of each method ^62,63^. We recognize that our combined map of watersheds identified with wild pigs represents a minimum estimate and that there are likely some undetected watersheds, but given such rapid expansion of wild pigs this is inevitable. Non-detection was addressed in the survey design and implementation with the use of multiple survey methods across the study area and a large sample size^62,63^. The possibility of false positives by misidentifying species is also a concern in presence/absence surveys that has the potential to overestimate presence^63^, however, wild pigs are a novel, large, and distinctively shaped species on the landscape that are unlikely to be misidentified. Most of the individuals who observed wild pigs live and work outdoors daily and have a much higher than average exposure to wildlife and have much experience and observations to share.

The overall goal of our comparison of different methods was to promote consideration of a wide range of options and how to evaluate and integrate them, not to be prescriptive. Each study will have different budgets, time constraints, and access to existing datasets and so will make different decisions. The use of GPS collars is a common technique in wildlife research, providing multiple locations daily with high spatial accuracy^64^, however they were by far the most expensive method evaluated and are generally best suited for more final scale analyses. Expert elicitation through systematic sampling provided the largest number of independent, unique locations at moderate cost. The use of expert elicitation is a common method used in modeling species distribution and is a valuable resource^65,66^, but is potentially limited by non-response bias. Non-response and self-selection bias, inherent to survey data collection, is reduced by the large sample size generated from multiple techniques, which allows for greater statistical power^67,68^. Access to social networks of stakeholders using snowball sampling provided the greatest number of independent locations across a broad spatial scale at a relatively low cost. The use of social networks allows for large sample sizes to be obtained at a low-cost from hard-to-reach populations with unknown parameters^69,70^, but it can introduce selection bias since this involves non-probability sampling of a specific target group, where over or under-representation of a group or specific characteristic can occur due to stronger or weaker social connections^71^. Sampling bias was addressed through the large sample of participants as well as by access to several small, discrete referral chains, rather than a few large ones^72,73^. The telephone survey was only moderately effective in terms of the number and range of wild pig detections compared to the high cost. However, the random stratified telephone sampling method and large sample size across all ten Canadian provinces provided some new data and helped examine if there was any bias in data obtained from the other social science methods. The Alberta bounty program provided a large number of wild pig locations relative to its small spatial scale but was limited to a single province. While bounties are considered counter-productive to wild pig control efforts^74^ they did help identify the provincial distribution. Research and citizen science trail camera photos provided a means of validation for wild pig sightings, therefore were a low-cost, spatially accurate, and unbiased means of validating locations. Media searches provided very little information, therefore are not recommended for future study designs.

The combined use of multiple methods that incorporates social and ecological data has been documented in the literature and is an effective tool for large-scale data collection of a species^75,32,34,33,76,77^. Large sample sizes, several, discrete referral chains, and the incorporation of multiple sampling methods conducted in this study provided a more robust dataset, captured a wider range of information holders, and reduced sampling bias ^76,77,78^. As such, we did not choose one method as “truth” against which to compare all others, but rather to consider all methods as any other dataset to evaluate benefits and limitations and compare with the database of all combined occurrences^79^. Several studies have found that a strong correlation exists between traditional or local ecological knowledge and western science, with the differences between the two ultimately stemming from differences in temporal and spatial scale ^33,32,80,81^. When used in conjunction with one another the variation in spatial and temporal scales between local and traditional knowledge with conventional ecological research provides complimentary and novel information and can address gaps in knowledge ^32,82,83^.

The overall approach that we present here can be applied to any invasive species, from plants to animals. The rapid identification of invasive species distribution provides managers with the ability to implement management strategies to attempt to eradicate the species before establishment occurs and the cost of eradication becomes unfeasible. In the current millennium with ever expanding globalization, land use changes, and climate change, challenges associated with invasive species is likely to only increase ^84,3^. Future research can apply the technique established in this study for rapid and cost-effective identification and understanding of the distribution of an invasive species on the landscape.

## METHODS

### Study Area

Our study area included all ten Canadian provinces. Bordered by the Pacific Ocean to the west and the Atlantic Ocean to the east, Canada’s provincial landmass has an area of 5,499,918 km^2 85^ and stretches from the U.S/Canada border northward to 60°00’ N latitude. This large expanse of land includes a broad range of biodiversity and variability in ecosystems, topography, temperature, and precipitation that includes mountain ranges, open grassland, temperate rainforest, coniferous and deciduous forest, plains, and boreal shield^86^. As a result of the heterogeneity across latitude, elevation, and proximity to the moderating effects of large waterbodies, or the more extreme inland climates, considerable variability in temperature and precipitation is observed across Canada’s 15 ecozones^87,88^.

Our preliminary research based on interviews with researchers and wildlife managers across Northern Canada’s three territories (Yukon, Northwest Territories, and Nunavut) who further consulted with field staff and stakeholders, determined that there was no evidence of wild pigs in these most northerly ecozones, as predicted based on the literature, the extreme cold winter temperatures (< −50°C) combined with long winters (>7 months), and the absence of past or present domestic wild boar production (except several recent small operations near Whitehorse, Yukon) (Brook unpublished data) which are the source of invasive wild pigs. As such we excluded Taiga Cordillera, Taiga Plans, Taiga Shield, Hudson Plains, Southern Arctic, Northern Arctic, Arctic Cordillera, and Tundra Cordillera Ecozones and defined our current study area as including the Pacific Maritime, Montane Cordillera, Boreal Plains, Boreal Shield, Prairies, Mixedwood Plains, and Atlantic Maritime Ecozones.

### Data Collection Comparisons

We collected eight unique, independent datasets using social science and conventional wildlife monitoring to document wild pig occurrences across Canada. Our social survey design was approved by the University of Saskatchewan Behavioral Research Ethics Board (BEH# 15-155) and the ecological data was approved by University of Saskatchewan Animal Research Ethics Board (Animal Use Protocol no. 20150024). While the term ‘wild pig’ is now widely used^6^, for our survey we used the term ‘wild boar’ as we found from preliminary consultations and pretesting that this term was best understood at the time of the study by rural Canadians. We are now consistently using ‘wild pigs’, recognizing that few, if any, pure bred domestic or free-ranging Eurasian wild boar exist in Canada. The datasets obtained were evaluated against one another to assess the trade-offs that occur from different data collection methods. The total number of wild pig detections and number of occupied watersheds were calculated along with the total cost and cost per wild pig detection for each data collection method (Table 3).

### Expert Elicitation Survey Design

A stratified sampling design based on provincial wildlife enforcement and management districts was conducted from 2015-2017. At least one Conservation Officer (CO) and one government wildlife biologist in each district was contacted where possible. Contact information and locations were found through employee directories on provincial government websites. The first positive response received was the individual with which the interview was completed. Three contact attempts were made for each individual, with a message left on the first attempt.
Attempts were halted once one interview within the strata had been completed or there was no response after three attempts. If no interviews were completed within a stratum it was considered a non-response. Strata were considered half-complete if both a CO and biologist were located in the stratum, but only one of the two completed an interview. Interview questions are included in Supplementary Methods (S1).

### Stakeholder Interview Snowball Sampling Design

A snowball sampling survey design^89^ was implemented to access key participants with knowledge of wild pig presence. Sampling took place from October 2014 to December 2017. Recruitment for this study began with an individual who shared their observations and then provided additional contacts that were also likely to know of wild pig occurrences. This snowball method of accessing additional contacts creates an ever-expanding network of individuals with information^77,90^. The technique uses social networks, therefore personal communication between the researchers and members of the social networks was essential^71^. Non-probability sampling methods typically require the use of advertising and outreach to be successful, compared to the stricter framework of probability sampling, in which letters and phone calls are made to specific individuals^91^. The snowball sampling method for this study used both techniques for recruiting participants. Multiple methods of recruitment were provided to gain access to participants. Presentations at wildlife meetings and conventions, information booths, posters, and magazine articles were all forms that were used to recruit participants. Active recruitment such as presentations and information booths were used to make initial contacts and gain access to social networks. Face-to-face contact was primarily used for initial contact and telephone was used to access additional contacts. Passive recruitment in the form of posters and magazine articles were used in an attempt to access an even broader geographic range of participants. Individuals with first-hand wild pig knowledge were also provided by the sample of experts. In order to keep the interest of the participant, and hence, response rate high, interviews were not completed with snowball sample members, only questions relating to wild pig locations were asked.

### Rural Telephone Survey Design

We used a stratified random sampling design to conduct a telephone survey of 3,000 individuals, representative of the rural Canadian population. The survey instrument included several themes but for the purposes of this paper we only used the responses to three questions: (1) What is your postal code? (2) Are you aware of feral boar presence or observations in your area in the past five years? (3) If yes, can you provide a date and location of the observation? Data collection was conducted by the Social Science Research Laboratory call center at the University of Saskatchewan. Participants could respond in either official language, English or French, and French responses were translated by individuals with a strong proficiency in that language. Provinces were stratified based on Forward Sortation Area’s (FSA’s) for rural areas as defined by Canada Post. The interview quota for each province was stratified based on the provincial rural population (from Stats Canada 2017 unpublished data) and individual contact information was randomly selected from within the FSA’s. Interviewers asked to speak with a member of the household that was over the age of 18 and was having the next birthday. Interviews were completed in February and March of 2016 to take advantage of the time when farmers were least busy. Individuals with wild pig information were asked if they would be willing to provide their contact information and receive a follow-up call from the primary researcher. Follow-up calls inquired about wild pig locations in greater detail as a means of validation. Interview questions and scripts are included in Supplementary Methods (S1).

### Bounty Data Collection

A bounty on wild pigs was in effect in the province of Alberta from 2003 to 2016. The program was initiated by the provincial government with individual counties signing up to participate. Hunters received $50 for a pair of wild pig ears and information regarding the location and date of the kill was turned in to the county. All counties participating in the program were contacted and data regarding the location, date, and number of pairs of ears turned in was received. Data were then compiled and verified.

### GPS Collars

We captured 12 female and 10 male feral wild boar by net-gun fired from a helicopter (n = 17) and through trapping using corral-style traps (n = 5) in 2015 and 2016. After capture, each feral wild boar was physically immobilized and fitted with a Global Positioning System (GPS) tracking collar (Telonics, Mesa, Arizona, USA). All collars were programmed to record a location every three hours and transmit the data via Iridium satellite link.

### Wild Pig Occurrence Mapping

All wild pig occurrences were converted into UTM coordinates using Google Maps (2018) and the Legal Land Description Converter (2017). Estimates of spatial accuracy of the data were based on information from the provider and using map scale. Each occurrence was buffered by 10 km^33^, a conservative estimate of annual home range movements and an area larger than the expected spatial error from any of the data collection methods. Two Level 9 (North America and Arctic) watershed shapefiles were downloaded from HydroSHEDS (2017) following the Pfafstetter coding system^92^. The 37,578 Level 9 watershed sub-basins in the study area have a mean area of 267 km^2 93^. Watershed units were chosen to model wild pig distribution as they are ecologically stratified units and allow for consistency across the landscape based on abiotic and biotic factors^10,33,94^. Level 9 watershed sub-basins are large enough to include the home range of at least one individual or sounder, which average 3.6 km^2^ for sows and 4.91 km^2^ for boar^95,96,97,98^, while encompassing considerable landscape heterogeneity^10^. The use of watershed units also maintains confidentiality of exact locations to ensure private land is not easily accessible to hunters, which was a condition of many landowners for providing information. Watersheds that intersected a wild pig location were selected to create a watershed occurrence layer illustrating wild pig presence by watershed units^33^. Two ten-year interval maps were created from 1990-2000 and 2001-2011. A current wild pig location map was created for the years 2012-2017.

## DATA AVAILABILITY

Data compiled and analyzed in regards to this study is available upon request from the corresponding author.

## ACKNOWLEDGEMENTS

This study was made possible through funding provided by the United States Animal and Plant Health Inspection Service National Feral Swine Damage Management Program, the University of Saskatchewan, Saskatchewan Fish and Wildlife Development Fund, Natural Science and Engineering Research Council (NSERC), and the Cyril Capling Trust at the University of Saskatchewan. This study would not be possible without the help of Raea Gooding, Chantal Jantzen, Dallis Aiken, Annie Rosia, Phillipa Williams, and Marissa Berard, as well as the generous contributions of our study participants.

